# Minimal effective dose of oral bedaquiline and activity of a long acting formulation of bedaquiline in the murine model of leprosy Bedaquiline for the treatment of leprosy

**DOI:** 10.1101/2023.05.15.540907

**Authors:** Aurélie Chauffour, Nacer Lounis, Koen Andries, Vincent Jarlier, Nicolas Veziris, Alexandra Aubry

## Abstract

**Background:** Bedaquiline (BDQ), by targeting the electron transport chain and having a long half-life, is a good candidate to simplify leprosy treatment. Our objectives were to (i) determine the minimal effective dose (MED) of oral BDQ, (ii) evaluate the benefit of adding another inhibitor of the respiratory chain to oral BDQ (*i.e.* clofazimine (CFZ)) and (iii) evaluate the benefit of an intramuscular injectable long-acting formulation of BDQ (intramuscular BDQ; BDQ-LA IM), in a murine model of leprosy.

**Methodology/ Principal Findings:** To determine the MED of oral BDQ and the benefit of adding CFZ, 100 four-week-old female nude mice were inoculated in the footpads with 5.10^3^ bacilli of *M. leprae* strain THAI53. Mice were randomly allocated into: 1 untreated group, 5 groups treated with oral BDQ (0.10 to 25 mg/kg), 3 groups treated with CFZ 20 mg/kg alone or combined with oral BDQ 0.10 or 0.33 mg/kg, and 1 group treated with rifampicin (RIF) 10 mg/kg. Mice were treated 5 days a week during 24 weeks.

To evaluate the benefit of the BDQ-LA IM, 340 four-week-old female Swiss mice were inoculated in the footpads with 5.10^3^ to 5.10^1^ bacilli (or 5.10^0^ for the untreated control group) of *M. leprae* strain THAI53. Mice were randomly allocated into the following 11 groups treated with a single dose (SD) or 3 doses (3D) 24h after the inoculation: 1 untreated group, 2 treated with RIF 10 mg/kg SD or 3D, 8 treated with oral BDQ or BDQ-LA IM 2 or 20 mg/kg, SD or 3D.

Twelve months later, mice were sacrificed and *M. leprae* bacilli enumerated in the footpad. All the footpads became negative with BDQ at 3.3 mg/kg. The MED of oral BDQ against *M. leprae* in this model is therefore 3.3 mg/kg. The addition of CFZ to a dose of BDQ 10-fold lower than this MED increased the bactericidal activity of BDQ suggesting synergies between both drugs. The BDQ-LA IM displayed similar or lower bactericidal activity than the oral BDQ.

**Conclusion:** We demonstrated that the MED of oral BDQ against *M. leprae* was 3.3 mg/kg in mice and the addition of CFZ to oral BDQ may improve the efficacy of BDQ. BDQ-LA IM was similar or less active than oral BDQ at equivalent dosing and frequency but should be tested at higher dosing in order to reach equivalent exposure in further experiments.

## Author summary

The current multidrug therapy is effective against leprosy but remains long and difficult to observe for patients supporting the need of monthly-based treatment. Bedaquiline (BDQ), a diarylquinoline with a long half-life, is a candidate drug to shorten leprosy treatment by targeting the electron transport chain and inhibiting the ATP synthesis. In this work, we demonstrated that (i) the minimal effective dose of oral BDQ against *M. leprae* is 3.3 mg/kg, (ii) the combination with another drug targeting the respiratory chain such as clofazimine may improve the efficacy of oral BDQ, and (iii) BDQ long acting formulation was similar or less active than oral BDQ at equivalent dosing and frequency but should be tested at higher dosing in order to reach equivalent exposure in further experiments.

## Introduction

Leprosy remains a major health problem worldwide despite being one of the oldest infectious diseases, reported for more than 2000 years. The leprosy elimination goal as a public health problem set by the World Health Organization, aiming for a global prevalence rate of < 1 patient in a population of 10,000, was achieved in 2000, but up to 200,000 new cases are still reported each year [1]. The worldwide use of leprosy drugs starting in the 1980s and their access at no cost for patients since 1995 were tremendous in the ability to achieve leprosy elimination [2]. The next goal which remains to achieve is to develop a strategy focusing on zero leprosy by the end of 2030 [3]. Several parameters impair this latter goal: (i) the reassignment of health activities due to the Covid-19 pandemic that may lead to a decrease in leprosy diagnosis and treatment and an underestimation of the number of leprosy cases [4], (ii) as with other bacteria of medical interest, antimicrobial resistance is observed in the causative agent *Mycobacterium leprae* in several parts of the world, despite multidrug therapy being the recommended standard leprosy treatment to avoid resistance selection since 1982. The first treatment of leprosy, consisting of a monotherapy of dapsone, led to the emergence of drug-resistance [5]. Despite the addition of rifampicin (RIF) in the 1960s, drug-resistant strains quickly emerged [6]. Moreover, the length of the treatment leads to a poor compliance by patients and may favor the emergence of resistant strains. Therefore, to simplify and to facilitate the direct observation of treatment, a shorter, fully supervisable, monthly-administered multidrug regimen for leprosy is highly desirable [7]. Finally, in addition to patients whose *M. leprae* isolates are resistant to RIF, special regimens are also required for individual patients who cannot take RIF because of allergy or intercurrent disease such as chronic hepatitis.

In 2005, a newly discovered class of antibiotics, the diarylquinoline, was reported to be highly bactericidal against *M. tuberculosis* in mice and later in the mouse models for *M. leprae* [7] [8] [9]. The lead compound, bedaquiline (BDQ), also called R207910 or TMC207, inhibits an enzyme belonging to the electron transport chain, the ATP synthase, by binding to the subunit c of the enzyme, leading to a decrease in bacterial metabolism. The bactericidal activity of oral BDQ against *M. leprae* observed in mice is similar to that of moxifloxacin and RIF supporting the launch of a clinical trial aiming at evaluating BDQ efficacy in multi bacillary (MB) leprosy [10]. Oral BDQ is currently the only new drug under clinical trial for leprosy treatment [10].

Interestingly, *M. leprae* does not possess all the proteins along the electron transport chain [11] suggesting that associating inhibitors acting at different enzymes belonging to it may act synergistically and may display strong bactericidal anti-leprosy activity. Clofazimine (CFZ), whose mechanism of action is not fully understood, targets the electron transport chain at the level of the menaquinone. CFZ has been shown to be effective against leprosy [12–20]. The anti-leprosy activity of the association of the two drugs acting on electron transport chain, e.g. BDQ and CFZ, deserve to be evaluated. In addition, the *Rv6708* gene that encodes for an inhibitor of efflux pumps in *Mycobacterium tuberculosis* is a pseudogene in *Mycobacterium leprae* where it might contribute to a higher potency of bedaquiline against *M. leprae* compared to *M. tuberculosis* [11].

Two key properties of drugs administered in LAI (Long Acting Injectable) formulations are low aqueous solubility to preclude the rapid dissolution and release of the active drug substance, and a reasonably long pharmacokinetic (PK) elimination half-life, *i. e.*, slow clearance from the body. For an antimicrobial, another desired property is high potency, negating the need for high concentrations in the blood and allowing lower drug doses to be injected. Bedaquiline seems to be highly potent against *M. leprae* [9] at a lower dose than against *M. tuberculosis*.

It has high lipophilicity (logP, 7.3), and a long half-life (about 24 h, functionally or effectively) which makes it suitable for use in an LAI formulation [21]. The efficacy of LAI BDQ has been already demonstrated in a latent tuberculosis infection mouse model [21–23]. Due to the very slow doubling time of *M. leprae*, a unique administration of standard of LAI BDQ could also be considered.

In our work, we aimed to (i) determine the minimum effective dose (MED) of the oral BDQ, (ii) evaluate the benefit of combining oral BDQ and CFZ, and (iii) evaluate the benefit of a BDQ long-acting formulation in a murine model of leprosy.

## Methods

### Ethics statement

The experimental project was favorably evaluated by the ethics committee n°005 Charles Darwin localized at the Pitié-Salpêtrière Hospital. Clearance was given by the French Ministry of Higher Education and Research under the number APAFIS#9575-2017030114543467 v3. Our animal facility received the authorization to carry out animal experiments (license number C 75-13-01) on the 27^th^ of April 2017. The persons who carried out the animal experiments had followed a specific training recognized by the French Ministry of Higher Education and Research and follow the European and the French recommendations on the continuous training.

### Materials

In both experiments, mice were infected with a *M. leprae* THAI53 strain. This strain was fully susceptible to the common antileprosy drugs (*i.e.* RIF, dapsone, CFZ and fluoroquinolones) [24]. The suspension used to inoculate mice was prepared from mice already infected by this isolate one year earlier. The Shepard and Mac Rae’s method was used to prepare the suspension [25]. Briefly, the tissue from the footpads were aseptically removed and grinded in a GentleMacs Octo Dissociator (Miltenyi®) under a volume of 2 ml of Hank’s balanced salt. Ten µl of the prepared suspension were taken to perform a Ziehl-Neelsen staining to count *M. leprae* Acid Fast Bacilli (AFB). Suspensions needed to inoculate mice were then further diluted in Hank’s balanced salt.

Respectively four-week-old nude (NMRI-*Foxn1*^nu/nu^) for the determination of the minimal effective dose of oral BDQ and Swiss mice for the evaluation of the BDQ-LA IM were purchased from Janvier Labs, Le Genest Saint Isle, France.

RIF and CFZ were purchased from Merck, France; oral BDQ and BDQ-LA IM were provided by Johnson and Johnson, Belgium.

### Infection of mice with *M. leprae* and treatment

First experiment: determining the minimal effective dose of oral BDQ against *M. leprae* and the contribution of CFZ when combined to BDQ. We adapted the continuous method to determine the MED of oral BDQ [26]. One hundred 4-week-old female nude mice were infected in the left hind footpad with 0.03 ml of the *M. leprae* isolate THAI53 according to the Shepard’s method [27] with an inoculum of 5.10^3^ AFB/ footpad. Mice were then randomly allocated into one untreated control group and 9 treated groups of 10 mice each: RIF 10 mg/kg, BDQ 0.10, 0.33, 1, 3.3 and 25 mg/kg, CFZ 20 mg/kg and combinations of BDQ 0.10 or 0.33 mg/kg and CFZ 20 mg/kg. Treatment was given one month after inoculation, five days a week during 24 weeks by oral gavage under a volume of 0.2 ml per mouse.

### Second experiment: comparing the bactericidal activity of oral BDQ and BDQ-LA IM against

***M. leprae***. We used the proportional bactericidal method that allows to measure the bactericidal activity of a compound [28]. Three hundred and forty 4-week-old female Swiss mice were infected in the left hind footpad with 0.03 ml of the *M. leprae* isolate according to the Shepard’s method [27]. Mice were inoculated with three different inocula of 5.10^3^, 5.10^2^, 5.10^1^ AFB/ footpad except for the untreated control which was also inoculated with one 5.10^0^ additional group. Mice were randomly allocated into one untreated control group and 10 treated groups of 10 mice each: RIF 10 mg/kg, oral BDQ 2 or 20 mg/kg, BDQ-LA IM 2 or 20 mg/kg. Treatment was given by oral gavage under a volume of 0.2 ml per mouse, except for the BDQ-LA IM which was injected intramuscularly under a volume of 0.012 ml per thigh and both thighs were injected at the same time. Treatment for all drugs was given as a single (SD) or three doses (3D) for BDQ 2 or 20 mg/kg and BDQ-LA IM 2 or 20 mg/kg and began the day after inoculation for the SD, and 4 and 8 weeks later for 3D.

### Assessment of the effectiveness of the treatment

To permit multiplication of *M. leprae* to a detectable level, mice were held 12 months in the animal facility. Mice were then euthanized and tissues from their footpad were removed aseptically and homogenized under a volume of 2 ml of Hank’s balanced salt according to the Shepard’s method [27]. *M. leprae* bacilli were considered to have multiplied (*i.e.* survived the treatment) in those footpads were found to contain ≥10^5^ acid-fast bacilli, regardless of the size of the inoculum.

### Statistical analysis

**First experiment: determine the MED of oral BDQ against *M. leprae***. A Mann-Whitney test was performed. A p-value <0.05 was considered to be statistically significant by standard evaluation. For multiple comparisons between the groups, Bonferroni’s correction was applied, *i.e.*, the difference would be significant at the 0.05 level only if the P value adjusted to the number of groups: 0.05/n in which n was defined as the number of primary comparisons. Thus, the corrected P was 0.05/10 = 0.005.

### Second experiment: compare the bactericidal activity of oral BDQ and BDQ-LA IM against

***M. leprae***. The proportion of viable *M. leprae* after treatment was determined as the 50% infectious dose. The significance of the differences between the groups was calculated by the Spearman and Kärber method [29]. A p-value <0.05 was considered statistically significant by standard evaluation. For multiple comparisons between the groups, Bonferroni’s correction was applied, *i.e.*, the difference would be significant at the 0.05 level only if the P value adjusted to the number of groups: 0.05/n in which n was defined as the number of primary comparisons. Thus, the corrected P was 0.05/11 = 0.0045.

## Results

### Minimal effective dose of oral BDQ (Table 1, Figure 1)

After one year of observation, all footpads were positive in the untreated control group (mean of 8.13±0.26 AFB per footpad), confirming the multiplication of *M. leprae*.

As compared to untreated control, RIF reduced the bacillary load by 5 log_10_ AFB (p=0.0002). The 3 lowest oral BDQ doses (0.10, 0.33 and 1 mg/kg) reduced the bacillary load by 1 log10 AFB as compared to untreated control (p=0.002, p=0.00001, p=0.0009 respectively), but mouse footpads remained positive. These 3 oral BDQ doses were also less bactericidal than RIF (p=0.0002 for the three groups). On the other hand, all the footpads were negative in the groups treated with the 2 highest oral BDQ doses (3.3 mg/kg and the 25 mg/kg) and displayed higher bactericidal activity than RIF 10 mg/kg (p≤0.006 for both groups, difference not significant after Bonferroni’s adjustment).

CFZ 20 mg/kg reduced the bacillary load by approximately 7 log_10_ AFB as compared to untreated control (p=0.0001) which is similar to RIF bactericidal activity (p=0.067, difference not significant after Bonferroni’s adjustment). The addition of CFZ to the 2 lowest oral BDQ doses (*i.e.* 0.10 or 0.33 mg/kg) lead to a decrease in the number of AFB positive footpads and reduced the bacillary load by approximately 6 log_10_ as compared with oral BDQ alone despite the reduction was not statistically significant (p=0.764 and p=0.386, respectively).

## Groups

### Comparison of oral BDQ and BDQ-LA IM (Table 2)

There were 2.75% viable *M. leprae* bacilli in the untreated group. The percentage of viable bacilli killed under treatment ranged between 74.86% and 99.85% depending on the treatment group.

Compared to untreated control, the percentage of viable bacilli was smaller in all groups except BDQ 2 mg/kg SD either oral or LA IM; but the difference between the untreated group and the RIF 10 mg/kg SD was not significant after Bonferroni’s adjustment. RIF 3D was significantly different from RIF SD (p<0.001).

When comparing groups treated with 2 mg/kg oral BDQ or BDQ-LA IM at the same frequency (SD or 3D), a similar number of footpads remained positive in both groups (p>0.05). When comparing groups treated with 20 mg/kg oral BDQ or BDQ-LA IM at the same frequency (SD or 3D), more footpads remained positive in the groups treated with BDQ-LA IM and the difference were statistically significant but not after Bonferroni adjustments whatever the number of administrations between BDQ and BDQ-LA IM groups (p>0.0045) (Table 2).

## Discussion

The treatment of leprosy remains a challenge due to the current length of the treatment. During the last decades, few new molecules were synthesized and even rarer are those active against *M. leprae*. In 2006, Ji et al. demonstrated the antileprosy activity of oral BDQ, a new diarylquinoline which was also active against other mycobacteria such as *M. tuberculosis* [7,8]. They showed that a single dose of 25 mg/kg BDQ was as effective as rifapentine, moxifloxacin, as well as RIF, which is currently the most powerful antileprosy drug. Nevertheless, the minimal effective dose of the oral BDQ is unknown despite knowing that it may enable developing a safe dose. In our present work, we determined that a dose as low as 3.3 mg/kg of oral BDQ displayed a bactericidal activity similar to a 25 mg/kg dose (Table 1, Figure 1). The MED of 3.3 mg/kg is higher than the ≤1 mg/kg that was found to be the lowest active dose in an immunocompetent mouse model of leprosy [9] which can be explained by the lack of T-cell immunity, known to be important in mycobacterial diseases, in the nude mouse model used to determine the MED of oral BDQ in the present study. The MED of oral BDQ was not achieved in the immunocompetent mouse model because all the doses tested including the lowest one (1 mg/kg) were able to reach the limit of detection of *M. leprae* present in mouse footpads as enumerated by the AFB microscopy [9] suggesting a MED of ≤ 1mg/kg. The MED of 3.3 mg/kg obtained in the immunocompromised mouse model of leprosy is lower than that obtained with *M. tuberculosis* in an immunocompetent mouse model (6.5 mg/kg) [8] but the MED of oral BDQ against *M. tuberculosis* should be compared to the MED of oral BDQ against *M. leprae* determined in an immunocompetent mouse model (MED ≤ 1 mg/kg) suggesting at least a 6-fold lower MED of oral BDQ against *M. leprae* compared to that of *M. tuberculosis*.

**Figure 1.**
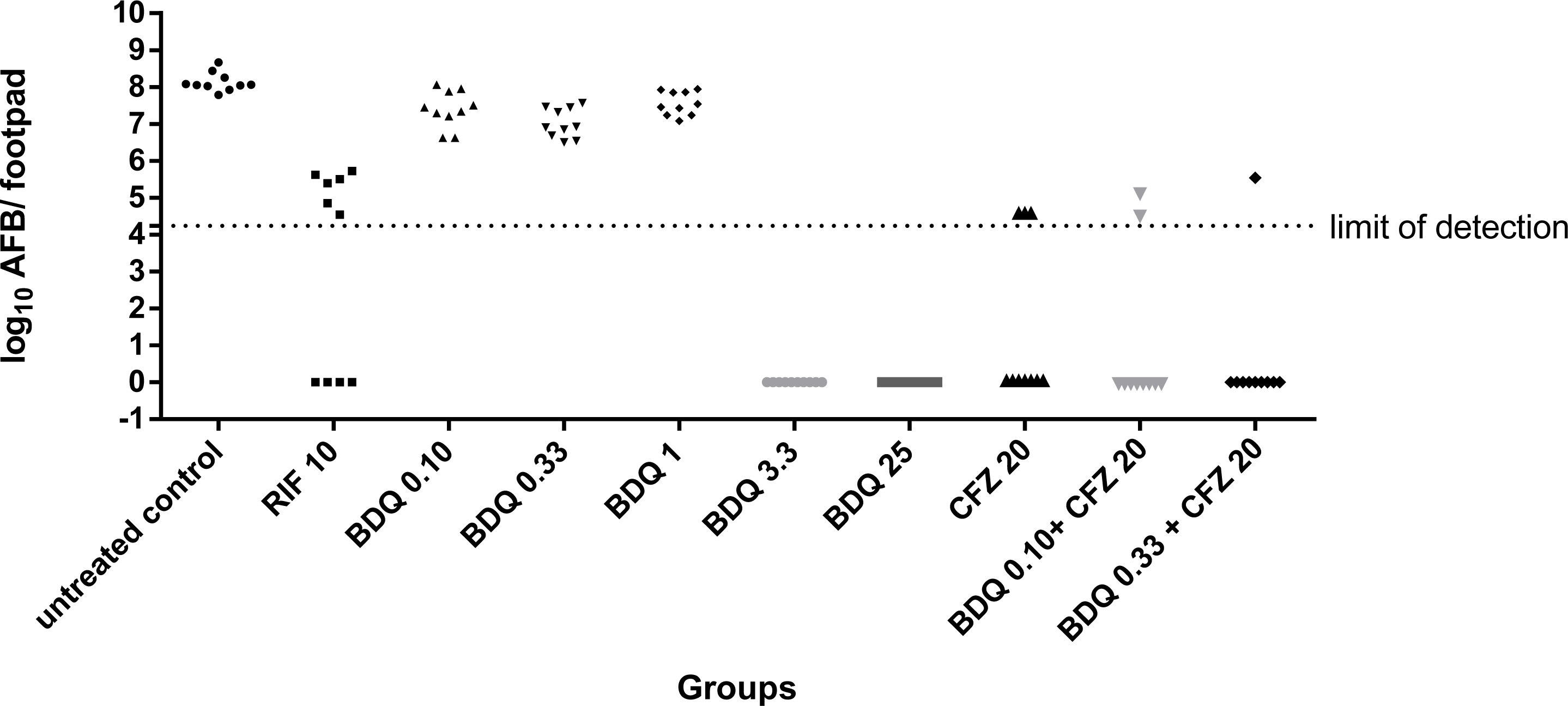
Multiplication of *M. leprae* organisms in mice treated by oral BDQ and the benefit of the combination of CFZ and BDQ (each mouse footpad is taken as a data point and the dotted line indicates the threshold of detection of *M. leprae*).

It’s important to mention that a single dose of 2 mg/kg oral BDQ (total dose = 2 mg/kg) in experiment 2 was able to kill 75% of the viable *M. leprae* bacilli while 6 months of 5 days per week treatment of 1 mg/kg (total dose = 120 mg/kg) in experiment 1 was not able to kill any bacilli present at start of treatment but was only able to slow down the replication of the bacilli when compared to untreated mice. The main differences between experiments 1 & 2 are the use of immunocompromised mice and initiation of treatment one-month post-infection in experiment 1 and the use of immunocompetent mice and initiation of treatment the day after infection in experiment 2. The use of immunocompromised mice without T cell immunity allows a much better replication of *M. leprae* and a much higher rate of viable bacilli while the replication rate and the rate of viable bacilli are much lower in immunocompetent mice which partially explain the differences in the effectiveness of the 2 regimens.

The treatment of leprosy needs to be based on a drug-combination to avoid the emergence of resistant-strains. BDQ targets the electron transport chain, which is also the target of the classical anti-leprosy drug CFZ [12,13]. The addition of CFZ to lower doses of BDQ which are not able to AFB convert mouse footpads allowed to reduce the number of positive footpads (Table 1). These 2 combinations were slightly more bactericidal than RIF. Despite being not statistically significant, this result may suggest that a combination of drugs targeting the electron transport chain may be highly bactericidal against leprosy. Further experiments should be designed and performed in nude mice rather than in Swiss, since the much larger numbers of *M. leprae* viable organisms in nude mice than in immunocompetent Swiss mice may permit more accurate differentiation among various levels of bactericidal activities.

**Table 1.** Multiplication of *M. leprae* organisms in nude mice to determine minimal effective dose of oral BDQ active against *M. leprae* and the benefit of the association of CFZ to BDQ.

| Treatment<sup>a</sup> | Positive footpad/<br>total footpad | Mean log<sub>10</sub> AFB/ footpad<br>median value (range of positive<br>footpads) | p value<sup>b</sup> |
| --- | --- | --- | --- |
| Untreated control | 10/10 | 8.13±0.26<br>(7.79-8.67) | / |
| RIF 10 mg/kg | 6/10 | 3.16±2.74<br>(4.54-5.72) | 0.0002 |
| BDQ 0.10 mg/kg | 10/10 | 7.39±0.50<br>(6.62-8.06) | 0.0019 |
| BDQ 0.33 mg/kg | 10/10 | 7.02±0.40<br>(6.50-7.57) | 0.00001 |
| BDQ 1 mg/kg | 10/10 | 7.56±0.32<br>(7.08-7.94) | 0.0009 |
| BDQ 3.3 mg/kg | 0/10 | ≤4.24 | ≤0.00006 |
| BDQ 25 mg/kg | 0/10 | ≤4.24 | ≤0.00006 |
| CFZ 20 mg/kg | 3/10 | 1.36±2.19<br>(4.54) | 0.0001 |
| BDQ 0.10 mg/kg + CFZ 20 mg/kg | 2/10 | 0.97±2.05<br>(4.54-5.15) | 0.0001 |
| BDQ 0.33 mg/kg + CFZ 20 mg/kg | 1/10 | 0.55<br>(5.54) | 0.00009 |

<sup>a</sup>, treatment was given by oral gavage 5 days a week during 24 weeks one month after inoculation,
<sup>b</sup>, comparisons versus untreated control (a p-value <0.005 was considered to be statistically significant when applying Bonferroni's correction).

One of the main characteristics of *M. leprae* is its long doubling time (*i.e.* 14 days) suggesting an active drug with a long half-life would be a good choice against this bacterium. The drug must be effective at a low dose and under an intermittent administration, conditions that are currently sought for the treatment of leprosy. A new formulation of BDQ, called BDQ-LA IM, was found to be active against latent tuberculosis [21]. Despite the potential challenges of introducing an injectable formulation in the field, its improved pharmacokinetics properties compared to oral BDQ may allow reduction in the duration of the treatment and therefore increase adherence of patients to the treatment. The BDQ-LA IM formulation was tested at two dosages, 2 or 20 mg/kg, and with two frequencies (one; SD or three doses; 3D). Our results showed that at equivalent dosing and frequency, oral BDQ was similar, or even more active than LA-IM in our murine model of leprosy (Table 2). A possible explanation is that the doses tested in our experiment were too low for LA-IM BDQ. Indeed, in the studies evaluating the BD-LA IM in tuberculosis higher doses were used and it was shown that BDQ LA-IM 160 mg/kg generated a C_max_ equivalent to that of 30 mg/kg oral BDQ. In support of this hypothesis is the fact that in the present study when comparing 20 mg/kg LA-IM BDQ to 2 mg/kg oral BDQ at equivalent dosing, the injectable form was more active than the oral form. Consequently BDQ LA-IM should be further tested at higher dosing against leprosy.

**Table 2.** Bactericidal activity against *M. leprae* THAI53 of oral bedaquiline and bedaquiline long-acting measured in Swiss mice by the proportional bactericidal method.

| Treatment<sup>a</sup> | No. of footpads showing multiplication<sup>b</sup> of *M. leprae*/<br>No. of footpads harvested, by inoculum |  |  |  | % viable *M. leprae*<sup>c</sup> | % viable *M. leprae* killed by treatment<sup>d</sup> | p value<sup>e</sup> | p value<sup>h</sup> |
| --- | --- | --- | --- | --- | --- | --- | --- | --- |
|  | 5.10<sup>3</sup> | 5.10<sup>2</sup> | 5.10<sup>1</sup> | 5.10<sup>0</sup> |  |  |  |  |
| Untreated control | 10/10 | 8/10 | 8/10 | 2/10 | 2.753 | / |  |  |
| RIF 10 mg/kg SD<sup>f</sup> | 8/10 | 6/10 | 4/10 | - | 0.432 | 90.01 | 0.018 | <0.001 |
| RIF 10 mg/kg 3D<sup>g</sup> | 0/10 | 0/10 | 0/10 | - | ≤0.004 | ≥99.85 | <0.001 |  |
| BDQ 2 mg/kg SD | 9/10 | 7/10 | 6/10 | - | 0.692 | 74.86 | 0.07 | 1 |
| BDQ-LA IM 2 mg/kg SD | 9/10 | 8/10 | 5/10 | - | 0.692 | 74.86 | 0.07 |  |
| BDQ 2 mg/kg 3D | 9/10 | 5/10 | 5/10 | - | 0.347 | 87.39 | 0.009 | 0.584 |
| BDQ-LA IM 2 mg/kg 3D | 8/10 | 6/10 | 3/10 | - | 0.219 | 92.04 | 0.0015 |  |
| BDQ 20 mg/kg SD | 3/10 | 0/10 | 0/10 | - | 0.009 | 99.67 | <0.001 | 0.005 |
| BDQ-LA IM 20 mg/kg SD | 7/10 | 3/0 | 1/10 | - | 0.055 | 98.00 | <0.001 |  |
| BDQ 20 mg/kg 3D | 0/10 | 0/10 | 0/10 | - | ≤0.004 | ≥99.85 | ≤0.001 | ≤0.014 |
| BDQ-LA IM 20 mg/kg 3D | 3/10 | 2/10 | 0/10 | - | 0.014 | 99.49 | <0.001 |  |

<sup>a</sup> treatment was given by oral gavage, except for BDQ-LA IM which was administered intramuscularly
<sup>b</sup> *M. leprae* bacilli were considered to have multiplied if the harvest from a footpad yielded >10<sup>5</sup> acid-fast bacilli
<sup>c</sup> the proportion of viable *M. leprae* after treatment was determined as the 50% infectious dose. The significance of the differences between the groups was calculated by the Spearman and Kärber method [29]
<sup>d</sup> calculated from the comparison of the proportion of viable organisms between untreated controls and the treated group.
<sup>e</sup> each treated group was compared to the untreated group of mice
<sup>f</sup> drug was given under a single dose the day after infection
<sup>g</sup> drug was given three times (the day after infection, 4 and 8 weeks later)
<sup>h</sup> p value corresponds to the comparison of the 2 groups at the beginning of the line: RIF SD vs 3D, BDQ oral vs LA-IM at equivalent dosing and frequency

In conclusion, we found that the MED of oral BDQ against *M. leprae* was 3.3 mg/kg and that the combination of clofazimine and BDQ may improve the bactericidal activity. BDQ-LA IM was similar or less active than oral BDQ at equivalent dosing and frequency but should be tested at higher dosing in order to reach equivalent exposure in further experiments. These findings open the path to the design of shorter BDQ-based treatment deserving to be evaluated in leprosy.

## References

1. Asia RO for S-E, Organization WH. Global leprosy strategy 2016-2020: accelerating towards a leprosy-free world - 2016 operational manual. WHO Regional Office for South-East Asia; 2016. Available: https://apps.who.int/iris/handle/10665/250119

2. WHO Study Group on Chemotherapy of Leprosy for Control Programmes, World Health Organization. Chemotherapy of leprosy for control programmes: report of a WHO study group [meeting held in Geneva from 12 to 16 October 1981]. 1982. Available: https://apps.who.int/iris/handle/10665/38984

3. World Health Organization. Towards zero leprosy. Global leprosy (Hansen’s Disease) strategy 2021–2030. World Health Organization; 2021. Available: https://apps.who.int/iris/handle/10665/340774

4. Global leprosy (Hansen disease) update, 2020: impact of COVID-19 on the global leprosy control. [cited 29 Nov 2021]. Available: https://www.who.int/publications-detail-redirect/who-wer9636-421-444

5. Pearson JM, Haile GS, Rees RJ. Primary dapsone-resistant leprosy. Lepr Rev. 1977;48: 129–132. doi:10.5935/0305-7518.19770016

6. Jacobson Robert R, Hastings Robert C. RIFAMPIN-RESISTANT LEPROSY. The Lancet. 1976;308: 1304–1305. doi:10.1016/S0140-6736(76)92071-7

7. Ji B, Chauffour A, Andries K, Jarlier V. Bactericidal activities of R207910 and other newer antimicrobial agents against Mycobacterium leprae in mice. Antimicrob Agents Chemother. 2006;50: 1558–1560. doi:10.1128/AAC.50.4.1558-1560.2006

8. Andries K, Verhasselt P, Guillemont J, Göhlmann HWH, Neefs J-M, Winkler H, et al. A diarylquinoline drug active on the ATP synthase of Mycobacterium tuberculosis. Science. 2005;307: 223–227. doi:10.1126/science.1106753

9. Gelber R, Andries K, Paredes RMD, Andaya CES, Burgos J. The Diarylquinoline R207910 Is Bactericidal against Mycobacterium leprae in Mice at Low Dose and Administered Intermittently. Antimicrobial Agents and Chemotherapy. 2009;53: 3989–3991. doi:10.1128/AAC.00722-09

10. Janssen Research & Development, LLC. An Open-Label Study to Evaluate the Efficacy and Safety of TMC207 in Subjects With Multibacillary Leprosy. clinicaltrials.gov; 2022 May. Report No.: NCT03384641. Available: https://clinicaltrials.gov/ct2/show/NCT03384641

11. Cole ST, Eiglmeier K, Parkhill J, James KD, Thomson NR, Wheeler PR, et al. Massive gene decay in the leprosy bacillus. Nature. 2001;409: 1007–1011.

12. Yawalkar SJ, Vischer W. Lamprene (clofazimine) in leprosy. Basic information. Lepr Rev. 1979;50: 135–144.

13. Black PA, Warren RM, Louw GE, van Helden PD, Victor TC, Kana BD. Energy Metabolism and Drug Efflux in Mycobacterium tuberculosis. Antimicrob Agents Chemother. 2014;58: 2491–2503. doi:10.1128/AAC.02293-13

14. Ji B, Perani EG, Petinom C, Grosset JH. Bactericidal activities of combinations of new drugs against Mycobacterium leprae in nude mice. Antimicrob Agents Chemother. 1996;40: 393–399. doi:10.1128/AAC.40.2.393

15. Ji B, Perani EG, Petinom C, N’Deli L, Grosset JH. Clinical trial of ofloxacin alone and in combination with dapsone plus clofazimine for treatment of lepromatous leprosy. Antimicrobial Agents and Chemotherapy. 1994;38: 662–667. doi:10.1128/AAC.38.4.662

16. Gelber RH. The killing of Mycobacterium leprae in mice by various dietary concentrations of clofazimine and ethionamide. Lepr Rev. 1987;58: 407–411.

17. Levy L. Activity of four clofazimine analogues against Mycobacterium leprae. Lepr Rev. 1981;52: 23–26.

18. Holmes IB, Banerjee DK, Hilson GR. Effect of rifampin, clofazimine, and B1912 on the viability of Mycobacterium leprae in established mouse footpad infection. Proc Soc Exp Biol Med. 1976;151: 637–641. doi:10.3181/00379727-151-39276

19. Shepard CC. Combinations involving dapsone, rifampin, clofazimine, and ethionamide in the treatment of M. leprae infections in mice. Int J Lepr Other Mycobact Dis. 1976;44: 135– 139.

20. Shepard CC. Minimal effective dosages in mice of clofazimine (B,663) and of ethionamide against Mycobacterium leprae. Proc Soc Exp Biol Med. 1969;132: 120–124. doi:10.3181/00379727-132-34162

21. Kaushik A, Ammerman NC, Tyagi S, Saini V, Vervoort I, Lachau-Durand S, et al. Activity of a Long-Acting Injectable Bedaquiline Formulation in a Paucibacillary Mouse Model of Latent Tuberculosis Infection. Antimicrob Agents Chemother. 2019;63: e00007–19. doi:10.1128/AAC.00007-19

22. Kaushik A, Ammerman NC, Tasneen R, Lachau-Durand S, Andries K, Nuermberger E. Efficacy of Long-Acting Bedaquiline Regimens in a Mouse Model of Tuberculosis Preventive Therapy. Am J Respir Crit Care Med. 2022;205: 570–579. doi:10.1164/rccm.202012-4541OC

23. Nguyen V, Bevernage J, Darville N, Tistaert C, Van Bocxlaer J, Rossenu S, et al. Linking In Vitro Intrinsic Dissolution Rate and Thermodynamic Solubility with Pharmacokinetic Profiles of Bedaquiline Long-Acting Aqueous Microsuspensions in Rats. Mol Pharm. 2021;18: 952–965. doi:10.1021/acs.molpharmaceut.0c00948

24. Matsuoka M. The history of Mycobacterium leprae Thai-53 strain. Lepr Rev. 2010;81: 137.

25. Shepard CC, McRae DH. A method for counting acid-fast bacteria. Int J Lepr Other Mycobact Dis. 1968;36: 78–82.

26. Shepard CC, Chang YT. Effect of Several Anti-Leprosy Drugs on Multiplication of Human Leprosy Bacilli in Foot-Pads of Mice. Proceedings of the Society for Experimental Biology and Medicine. 1962;109: 636–638. doi:10.3181/00379727-109-27293

27. Shepard CC. The experimental disease that follows the injection of human leprosy bacilli into foot-pads of mice. The Journal of experimental medicine. 1960;112: 445–454.

28. Colston MJ, Hilson GR, Banerjee DK. The “proportional bactericidal test”: a method for assessing bactericidal activity in drugs against Mycobacterium leprae in mice. Lepr Rev. 1978;49: 7–15.

29. Shepard CC, others. Statistical analysis of results obtained by two methods for testing drug activity against Mycobacterium leprae. Int J Lepr Other Mycobact Dis. 1982;50: 96–101.

